# Artificial Recombination of Nonribosomal Peptide Synthetases Rapidly Evolves Natural Products

**DOI:** 10.64898/2026.02.09.704528

**Authors:** Lin Zhong, Xingyan Wang, Xingxing Shi, Xianping Bai, Yang Liu, Hanna Chen, Boopathi Seenivasan, Xiaoqi Ji, Qingsheng Yang, Shengying Li, Rolf Müller, Qiang Tu, Youming Zhang, Xiaoying Bian

## Abstract

Efficient engineering of nonribosomal peptide synthetases (NRPSs) is a key strategy for expanding valuable peptide natural products. Current bioinformatics-guided approaches are often constrained by a set of predefined fusion sites, remaining experimentally challenging in most NRPS systems. Here we present the Recombineering Accelerated Evolution (RACE), which harnesses Red/ET recombineering mediated partially matched homologous recombination to recapitulate recombination-driven NRPS evolution on a highly accelerated timescale. Application of RACE to six known NRPS gene clusters generated 830 recombinants and yielded over 600 novel peptides including novel bis-lipopeptides. These recombinants reveal 112 previously unrecognized recombination fusion sites, providing an extensive landscape for NRPS evolution and more useful resources for guiding NRPS engineering. The RACE establishes a new paradigm for programmable NRPS evolution and enables rapid discovery of bioactive peptides.

## Introduction

Natural products are vital sources of clinical drugs and key active components in agricultural biocontrol agents [1-3]. Nonribosomal peptide (NRP) drugs, such as antibiotics like penicillin, vancomycin, polymyxin, daptomycin, the immunosuppressant cyclosporine, and the anticancer agent bleomycin, make up a significant fraction of natural product drugs [4]. NRP natural products are synthesized by a class of multi-modular enzymes called nonribosomal peptide synthetases (NRPSs) [5]. These multi-modular megasynthetases can be further divided into different domains, with three essential domains constituting a basic module: the adenylation (A) domain, responsible for activating and transferring one amino acid to the thiolation (T) domain, and the condensation (C) domain, which joins specific amino acids on the T domain to form the elongated nascent peptide chain. Multi-modular synthetases act as an “assembly line” for the synthesis of nonribosomal peptides, ultimately released by the thioesterase (TE) domain. The diversity of such natural products mostly depends on the A domain’s recognition of different amino acid substrates including an enormous variety of non-natural amino acids and post-assembly line modifications by different enzymes [6, 7].

The modular multienzyme NRPS machinery has generated significant structural and functional diversity of NRPs. But the total chemical synthesis of these complex and diverse natural products poses challenges[8]. Enhancing the pharmaceutical properties of natural products through semi-synthesis remains to be the prevailing approach in drug development[9, 10]. Genetic engineering of natural products provides a faster and more cost-effective way to unlock their pharmaceutical potential compared to traditional methods[11]. Efficient engineering of multi-modular systems like the NRPS has been a long-standing challenge and hot topic[12-14]. Due to the modular nature of NRPSs, where each module incorporates a specific building block, significant efforts have been devoted to engineering NRPSs to optimize their products. These strategies, collectively known as the combinatorial biosynthesis of NRPSs, have led to significant advances in modifying these megasynthetases to produce natural product analogues or even entirely novel, artificial compounds[13, 15-24]. The Bode group defined a fusion site (XU) between C and A domains, XUC fusion site between the N-lobe subdomain and C-lobe subdomain of C domain for NRPS refactoring[16, 24]. Calcott et al. successfully employed A-domain swapping as an effective strategy for NRPS engineering[21]. Although extensive efforts have been made to identify a universal fusion site for NRPS reprogramming[18, 19, 22, 23, 25-31], traditional trial-and-error strategies remain intrinsically limited by the extensive evolutionary diversity and structural complexity of NRPSs. Recent bioinformatic analyses have nevertheless revealed recombination as a major evolutionary driver of nonribosomal peptide (NRP) diversification[32-39], facilitating the identification of a limited number of previously undocumented fusion sites and improving engineering success in modular NRPS and polyketide synthase (PKS) systems[15, 40].However, engineering rules restricted to predefined fusion sites are fundamentally misaligned with the recombination-driven evolutionary logic of NRPSs, likely accounting for their persistent inability to reproduce the breadth, efficiency, and robustness of natural NRPS evolution. These limitations underscore the need for a new engineering paradigm that directly emulates recombination-driven evolution rather than imposing rigid, site-constrained designs (site-guided engineering strategies).

To achieve this goal, we developed a strategy that directly harnesses experimentally controllable recombination to drive NRPS evolution and engineering. Red/ET recombineering, which employs bacteriophage-derived recombinases Redαβγ/RecET, mediates both linear-plus-circular (LCHR) and linear-plus-linear (LLHR) homologous recombination via short homology arms (HAs) and has been widely established as a rapid, efficient, and user-friendly tool for genetic manipulation, including direct cloning, site-directed mutagenesis, and precise modification of NRPS and polyketide synthase (PKS) biosynthetic gene clusters[41-48], Notably, previous Red/ET-based NRPS mutagenesis[49] frequently produced truncated constructs lacking one or more modules due to recombination between homologous sequences of neighboring modules. This phenomenon is quite similar to the deletion event for diversification of NRPSs in natural evolution, providing the potential for experimental recapitulation of recombination-driven evolution of NRPSs. these deletion events mirror the module loss events that drive NRPS diversification in natural evolution, revealing the potential to experimentally recapitulate recombination-driven NRPS evolution. On the other hand, recombination-driven evolution in nature yields NRPS metabolites of exceptional structural diversity, reflecting abundant intrinsic recombination interfaces (homology arms) within NRPS gene clusters. These endogenous homology arms establish the molecular foundation for artificial NRPS evolution. Based on the Red/ET recombineering mediated partially matched homologous recombination, we established an accelerated recombination-derived evolution platform for NRPS diversification, termed <u>R</u>ecombineering <u>Ac</u>celerated <u>E</u>volution (RACE). RACE enables rapid NRPS evolution on a short timescale, supporting continuous evolution, *de novo* design, and rational engineering, ultimately enabling the custom synthesis of novel macrocyclic peptides and bioactive bis-lipopeptides.

## Results

### Experimental Mimicry of Natural Recombination Events by RACE

To mimic the natural recombination process, we introduced the concept of “Recombineering Accelerated Evolution (RACE)”, in which gene clusters are delivered into a heterologous host via DNA co-transformation to simulate HGT, and employs Red/ET recombineering to mediate recombination between designed donor fragments and recipient gene clusters. These yields recombinants carrying domain or module deletions, replacements, or insertions within a short timescale (Fig.1). Their sequences and products are then analyzed to directly link recombination events with product structures, recapitulating natural NRPS evolution process.

**Fig. 1.**
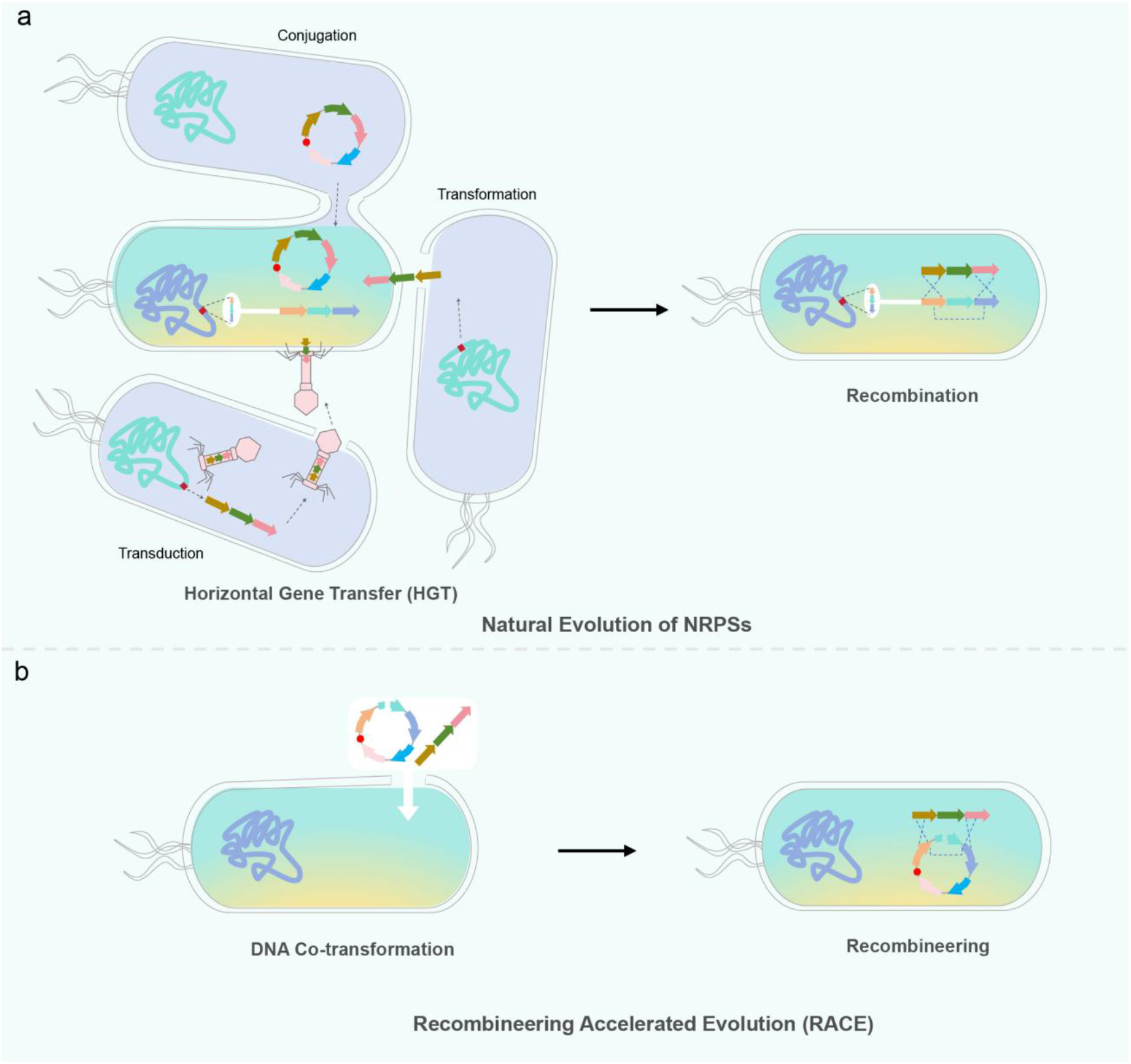
Recombineering Accelerated Evolution (RACE) technology mimics natural evolutionary processes in NRPSs. **a**. the natural evolution of NRPSs involves two primary mechanisms: horizontal gene transfer (HGT) and recombination. HGT includes transformation, transduction, and conjugation. **b**. RACE simulates these processes by co-transforming DNA to replicate HGT and utilizing rapid recombineering techniques to emulate natural recombination events, enabling accelerated NRPS evolution.

During the natural recombination-driven evolution of NRPSs, recombination HAs may be progressively diluted and probably occur within the short, discontinuous motifs distributed across the domains. However, the previous elaborate Red/ET recombineering usually requires HAs of ∼40 bp to achieve high-efficiency recombination[50, 51]. Thus, recapitulating these natural recombination events using Red/ET recombineering is only possible if this system can mediate efficient recombination under partial HA pairing. We adjusted the identities of the 40 bp downstream HA and varied the upstream HA to evaluate the effect of homologous pairing on recombination efficiency in the LLHR experiment (Fig. S1). Four groups were designed: 1) two 8 bp regions flanking a single mismatch (S2-S4), with S1 (perfect 8+1+8 bp) as positive and S5/S6 (single 8 bp match) as negative controls; 2) two 8 bp regions flanking two mismatches (S7, S8), with S7 as positive and S5/S6 as negative controls; 3) two 8 bp regions flanking three mismatches (S10); and 4) three 8 bp regions (upstream, central, downstream) containing two mismatches (S11, S12), with S11 as positive and S5/S6 as negative controls (Fig.S1). The results showed that all partially matched HAs mediated efficient RecET recombination, whereas the negative controls S5 and S6 (only 8 bp HA) yielded no correct recombinants (Fig. S2a, b). Constructs S2-S4 achieved 100% accuracy but showed a 10-to 14-fold lower efficiency than the fully matched control S1 (Fig. S2b). Similarly, S8, S10, and S12 produced correct recombinants (Fig. S2d), with S8 showing a 36-fold reduction relative to S7 and S10 a 175-fold reduction relative to S9 (Fig. S2c). In contrast, S12 exhibited only a ∼6-fold reduction compared with S11, indicating that additional short HAs mitigate the impact of mismatches. According to the strand annealing model of Red/ET recombineering, partially matched HAs (PMHAs) undergo RecT-mediated annealing to initiate recombination (Fig. S3) [44], with mismatches likely resolved by DNA repair, yielding a homogeneous recombinant. Notably, mismatched bases were preferentially corrected to the vector-derived sequence over the insert (Fig. S2b, d), possibly reflecting the bias of exonuclease activity during mismatch repair.

Compared with other *in vivo* recombination systems, e.g. CRISPR/Cas-induced approaches, this mechanism here offers distinct advantages by enabling efficient, precise homologous recombination using multiple discontinuous ultrashort paired regions as the HAs. Its efficiency is only marginally lower than that of fully paired homologous recombination, suggesting that the multiple discontinuous short homologous sequences (e.g. motifs) of NRPS gene clusters can be used as PMHAs for occurrence of the recombination within and across the NRPS gene clusters with the help of Red/ET recombineering. This provides an effective means to mimic natural recombination events of NRPSs, allowing us to establish a technological basis for accelerating NRPS evolution process by RACE.

We established RACE through two recombination modes: 1) intra-gene cluster evolution and 2) inter-gene cluster evolution (Fig. 2).

**Fig. 2.**
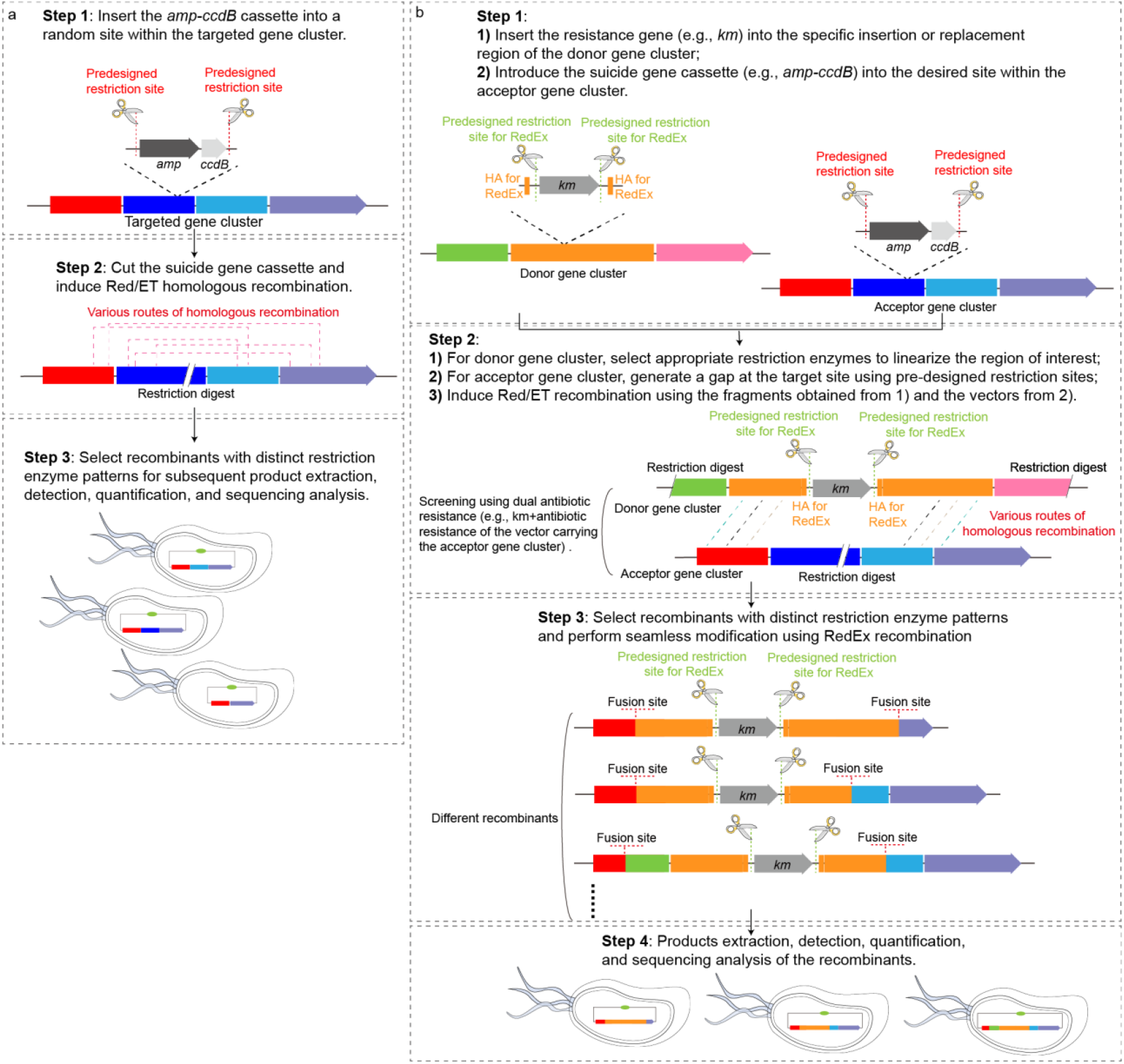
Recombineering ACcelerated Evolution (RACE). **a**. the intra-gene cluster evolution and **b**. inter-gene cluster evolution.

Intra-gene clusters evolution: The intra-cluster evolution strategy involved inserting antibiotic *resistance-ccdB* cassettes at various positions within gene clusters, followed by *in vitro* excision and RACE recombination (Fig. 2a and Fig. 3a). The *rzmA* (rhizomide) and *holA* (holrhizin) clusters (from *Mycetohabitans rhizoxinica* DSM 19002) and the *gdn* (glidonin) and *glp* (glidopeptin) clusters (from *Caldimonas brevitalea* DSM 7029) were used to implement intra-cluster evolution[18, 42, 52]. From *rzmA*, 41 unique recombinants yielded 20 classes of novel derivatives (**Ap3**-**Ap22**) (Fig. 3a-c, Figs. S4 to S7); *holA* generated 29 recombinants producing 13 classes of derivatives (**Ap24**-**Ap36**) (Fig. 3a-c, Fig. S4, Figs. S8 to S10); *gdn* produced 66 recombinants corresponding to 37 classes of novel products (**Ap38**-**Ap51, Ap53**-**Ap58, Ap60**-**Ap64, Ap66**-**Ap76, Ap78**) (Fig. 3a-c, Fig. S4, Figs. S11 to S12, Fig. S15); and *glp* yielded 82 recombinants resulting in 25 classes of new derivatives (**Ap81**-**Ap97, Ap99**-**Ap102, Ap104**-**Ap107**) (Fig. 3a-c, Fig. S4, Figs. S13 to S14, Fig. S16). Intra-cluster evolution produced remarkable product and recombination diversity, including not only single fusion (deletion) events but also multiple fusions and fusions with insertions (Fig. 3b), exemplified by recombinants A42/A43 (*rzmA*; Figs. S5, S6-4), A141/A142 (*gdn*; Figs. S11-3, S11-4), and A223-A226 (*glp*; Figs. S13-2, S13-3 and S13-5). Even a single generation of intra-cluster recombination generated complex derivative peptides through combinatorial reshuffling (Figs. S7-3, S15-32, S16-8), demonstrating the high-throughput potential of RACE. Novel product generation rates were high for *rzmA, holA*, and *gdn* (100%, 100%, and 90.9%, respectively), with median relative yields of 276%, 100%, and 99% compared to WT (Fig. S4). In contrast, *glp* showed a lower generation rate (59.7%) and a reduced median relative yield (3%) (Fig. S4 and Fig. 3b). These results suggest that NRPSs exhibit diverse evolutionary trajectories and substantial evolutionary plasticity, underscoring the variable complexity and challenges associated with their engineering.

**Fig. 3.**
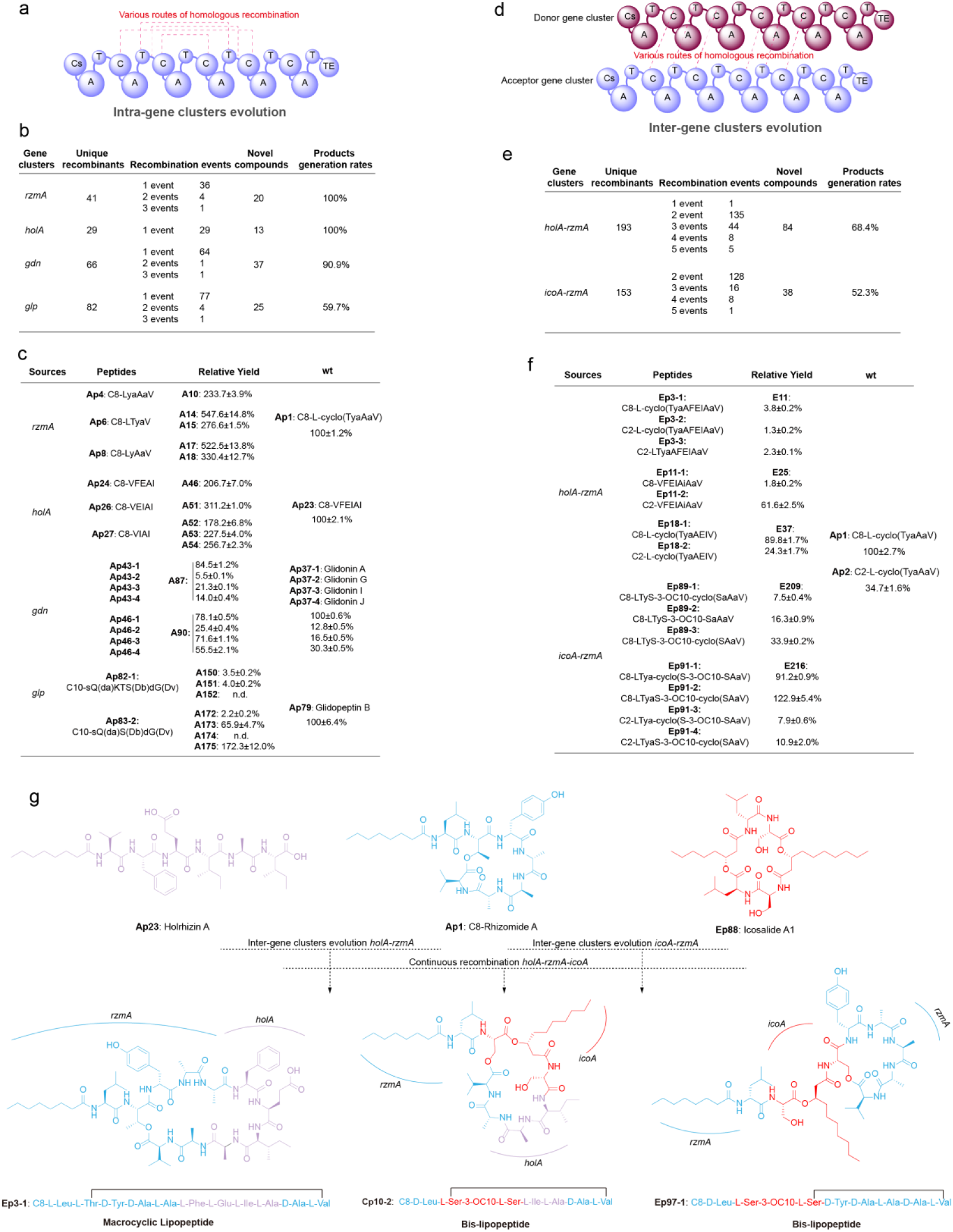
Statistical analysis of intra- and inter-gene cluster evolution. **a-c**. summary of intra-gene cluster evolution, including the number of recombinants and novel derivates, recombination events, and product generation rates (calculated as the number of product-forming recombinants divided by the total number of recombinants). Representative examples of newly derived products and their relative yields are also shown. **d-f**. corresponding statistical analyses for inter-gene cluster evolution. **g**. examples of lipopeptides obtained by RACE, including macrocyclic lipopeptides and the special bis-lipopeptides.

Inter-gene clusters evolution Inter-cluster RACE was performed in two designs (Fig. 2b, Fig. 3d): 1) the *rzmA* cluster served as the recipient with *holA* from the same strain as donor; 2) *rzmA* was again the recipient, with the *icoA* (icosalide) cluster from the distantly related strain *Burkholderia gladioli* ATCC 10248 as donor. The *icoA* cluster contains a unique C-Cs (starter condensation) didomain architecture that introduces a second lipid chain in its lipopeptide product. In the first design, an inducible *cm-tetR-tetO-ccdB* cassette was inserted into the *holA* A_4_ domain, and six defined gaps were introduced into *rzmA* (between the domains of Cs-A_1_, A_2_-T_2_, T_3_-Cdual_4_, T_4_-Cdual_5_, T_5_-C_6_, T_6_-Cdual_7_). This yielded 193 recombinants producing 84 classes of novel derivatives (**Ep1-Ep87**), with a median yield of 23% relative to WT (Fig. 3e, g, Figs. S17-S19, Fig. S22). On the one hand, naturally occurring bis-lipopeptides such as icosalides display potent antibacterial and antitumor activities[53]. On the other hand, introducing a second lipid chain into polymyxin through chemical synthesis markedly enhances its activity[54]. These findings collectively indicate that a second lipid chain can substantially strengthen lipopeptide bioactivity. Guided by this rationale, we sought to engineer artificial bis-lipopeptides in a high-throughput manner by exploiting the unique C-Cs domain of *icoA* through the RACE strategy. Therefore, in the second design, the cassette was inserted between the *icoA* C-Cs domains, generating 153 recombinants and 38 classes of novel products (**Ep89-Ep121, Ep123-Ep127**), with a median relative yield of 33% relative to WT (Figs. 3E, G, Fig. S17, Figs. S20-S21, and S23). Like intra-cluster evolution, inter-cluster evolution also produced extensive compound diversity. Notably, *rzmA-icoA* recombinants generated lipopeptides with two fatty acid chains (bis-lipopeptide), representing the first generation of bis-lipopeptides via combinatorial biosynthesis (Figs. 3f, g). Multiple recombination events were frequently observed, one-third of recombinants (E141-E201) carried more than two fusion sites in *rzmA-holA* recombinants, with groups such as E153, E155, E171, E183, and E192 containing up to five (Fig. 3e, Figs. S19-3, S19-4, S19-6, S19-17, S19-18). Similarly, in the *rzmA-icoA* recombinants, several (E330-E355) carried more than two fusion sites, with E351 harboring as many as five (Fig. 3e, Fig. S21). These findings indicate the high-throughput reshuffling capacity of RACE and the complexity of NRPS recombination preferences.

In total, 564 unique recombinants were obtained from intra- and inter-gene cluster evolution, yielding ∼218 classes of novel derivatives. Products included complex macrocyclic scaffolds and unprecedented structures with two fatty acid chains. The statistics of all recombinants and the relative quantification of products are presented in Figs. S5-S6 (for *rzmA*), S8-S9 (for *holA*), S11-S12 (for *gdn*), S13-S14 (for *glp*), S18-S19 (*rzmA* with *holA*), and S20-S21 (*rzmA* with *icoA*). These results demonstrate the high efficiency of RACE while emphasizing the intrinsic diversity of NRPS evolution and the difficulty of defining universally compatible recombination sites (Table S1).

#### Continuous evolution of NRPS

Continuous evolution of large modular enzymatic systems, such as NRPSs, remains a major challenge in the field of combinatorial biosynthesis. To evaluate the potential of RACE for driving continuous recombination, we performed a second round of recombination using first-generation recombinants as the starting acceptor (Fig. S24, Fig. 4a). The recombinase system was further optimized: the initial round employed both RecET and Redαβγ recombinases to facilitate LLHR and LCHR between the *rzmA* acceptor cluster and the *holA* donor cluster. Recombinants containing distinct fusion sites were pooled and used as acceptors for a second round of recombination. In this iteration, Redαβγ-mediated LCHR recombination was performed directly without prior linearization, using the *icoA* cluster as the donor (Fig. S24).

**Fig. 4.**
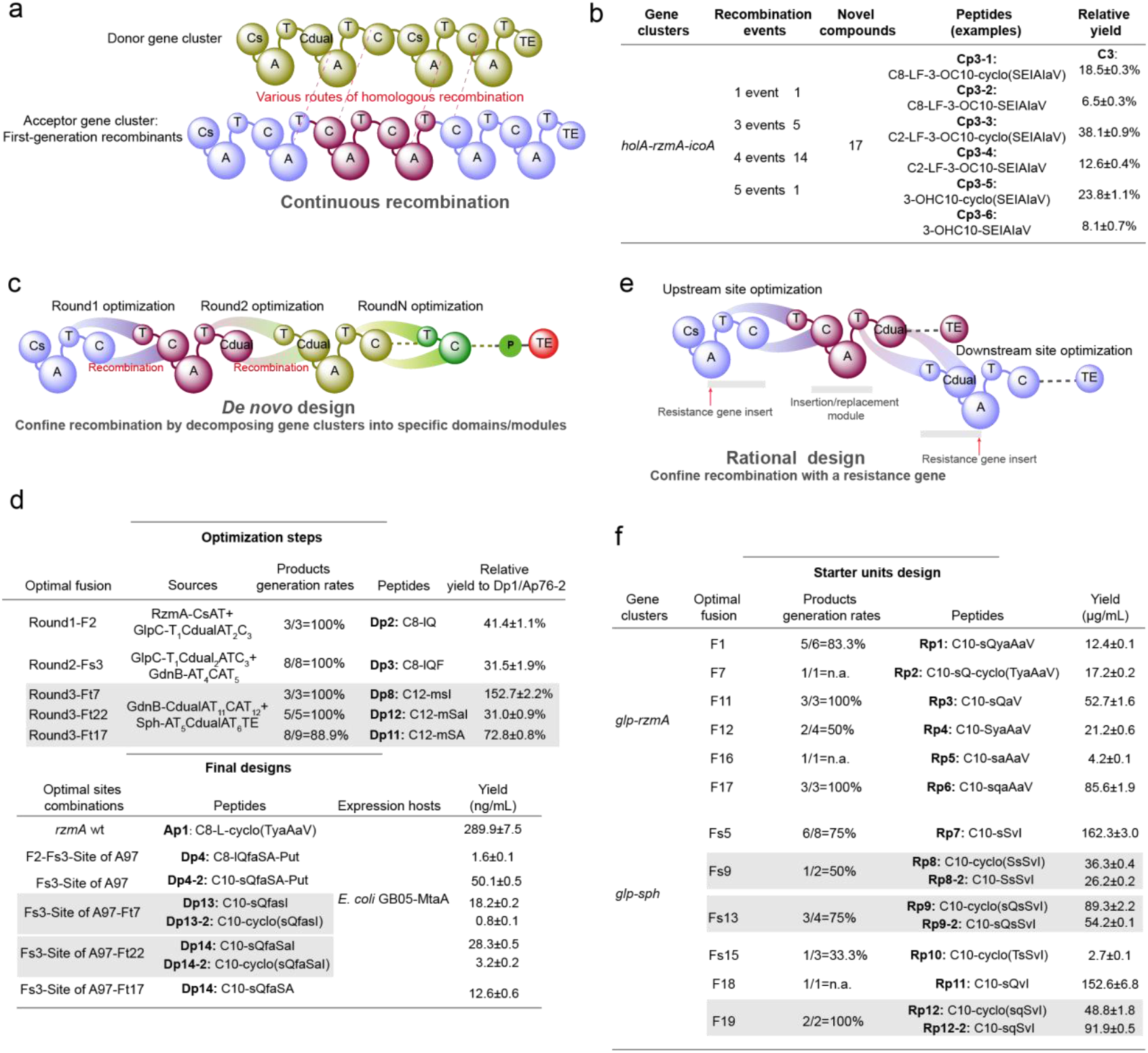
Statistical analysis of continuous recombination, *de novo* design, and rational design of the starter unit. **a-b**. summary of continuous recombination, showing the number of novel derivatives, recombination events, representative new products, and their relative yields. **c-d**. Summary of *de novo* design. This process consists of the optimization step followed by a final design step. The optimization focuses on selecting optimal fusion sites through recombination of split module combinations and recombinant quantification. The final step applies the selected sites to design the full-length gene cluster. **e-f**. rational design of the starter unit of lipopeptide products. In this design, recombination positions were unrestricted to maximize product diversity. It includes two parts: RACE-mediated recombination between the starter unit of the *glp* gene cluster and the main body of *rzmA*, and between the starter unit of *glp* and *sph*.

The successful generation of second-generation recombinants demonstrated the feasibility of RACE-enabled continuous recombination. Moreover, the use of Redαβγ-mediated LCHR significantly improved the efficiency of continuous recombination. To validate this approach, ten distinct first-generation recombinants were randomly pooled and recombined with *icoA* gene cluster via RACE. Sequencing of productive clones identified 21 second-generation recombinants (C1-C21), among which yielded 17 unique bis-lipopeptides (**Cp1**-**Cp17**), with a median yield of 26% relative to WT (Fig. 3g and Fig. 4a, b, Figs. S25-S27). The high titers of these novel derivatives show the power and scalability of continuous recombination via RACE (Fig. 4b, Figs. S25-3 to S25-5).

#### De novo design of artificial NRPs

RACE enables selection of optimal fusion sites based on the preferred recombination points as shown in intra and inter-gene clusters evolution (Figs. S5, S8, S11, S13, S18 and S20), making it suitable for *de novo* NRPS design. This strategy divides NRPSs into modules/domains to accelerate their evolution, with the thioesterase (TE) domain expressed independently to release intermediates (validated in preliminary experiments, Fig. S28). After each optimization cycle for maximal yield, evolution proceeds to the next round, ensuring optimal fusions at each site (Fig. S29).

The starter unit of *rzmA* gene cluster and GlpC-T_1_Cdual_2_A_2_T_2_C_3_ module of *glp* gene cluster were used as the first-round recombination of RACE. The terminal module (C_13_A*T_13_TE_1_TE_2_) of GdnB in *gdn* BGC was selected to release the product[18], because the mutant utilizing this module exhibited a higher product yield (Fig. S28). In the first round of RACE (Fig. S30a, c and Fig. S31), three different recombinants were obtained (Fig. 4c, d). The optimal fusion F2 (plasmid D8) yielded 41.4% production of **Dp2** compared to the production of **Dp1** in D10. Subsequently, F2 was utilized for the second round of RACE along with the GdnB-T_4_C_5_A_5_T_5_ of the *gdn* gene cluster to create the fusion Fs1-Fs8 (plasmids D12-D19) (Fig. 4d, Fig. S30b, c and Fig. S31). The optimal fusion, Fs2, resulted in a 31.5% production of **Dp3** compared to the production of **Dp2** in D8.

Finally, the fusion of F2 and Fs3, along with the fusion site from the truncated recombinants A97 as mentioned intra-gene clusters evolution (Fig. S14-4), was selected to design the final recombinant D23 (Fig. S30-S31). The final designed product **Dp4**, containing a unique C-terminal putrescine moiety, exhibited an absolute yield of 6.1 μg/L in *E. coli*. This yield is ∼60% lower than that of the A97 product (**Ap51-3**, 15.9 μg/L), likely due to cumulative deleterious effects from multiple site perturbations (Fig. 4d, Fig. S31d).

To generate novel cyclized derivatives, the initial D-Leu in **Dp4** was substituted with D-Ser by replacing the RzmA starter module with GlpC in recombinant D23, creating D24 (Fig. 4d, Fig. S32). A subsequent RACE round targeting the terminal module of D24, using the stephensiolide (*sph*) gene cluster of *Serratia marcescens* W1 and truncated recombinant A140 from the *gdn* cluster as the donor, yielded 26 recombinants. Six derivatives (**Dp7-Dp12**) were obtained (Fig. 4d, Figs. S33-S37). Among them, optimal fusions Ft7 (D35), Ft17 (D45), and Ft22 (D50) produced **Dp8, Dp11**, and **Dp12**, which were further used to design cyclized products in D24 (**Dp4-2**; Fig. 4d, Fig. S38). Final products included **Dp13/Dp13-2** and **Dp14/Dp14-2** (cyclized), as well as **Dp15** were designed, with absolute yields of 0.8, 18.2, 3.2, 28.3, and 12.6 μg/L in *E. coli* (Fig. 4d, Fig. S32 and Fig. S39). The reduced yields of cyclized products likely result from limited TE cyclization efficiency with unnatural D-Ser compared to native L-Thr, although partial cyclization with L-Ile was still observed (Fig. 4d, Fig. S33).

Collectively, the *de novo* design strategy achieved an overall product formation probability of 70%, with median relative yields of 20% and 52% for recombinants whose fusion sites were used to generate Dp4 and Dp7-Dp11, respectively. These results prove the high efficiency of the RACE method.

#### Rational modification of NRPS (Site-specific modification)

The first rational modification-RACE design for NRPS is to engineer the starter unit[55, 56], using the *glp* gene cluster’s starter unit to swap those of the *rzmA* and sph gene clusters. To maximize validation examples, recombination regions within the acceptor gene clusters (*rzmA* and *sph*) are not restricted (Fig. 4e, Fig. S40a and Fig. S41). This generated 39 recombinants, 19 with *rzmA-glp* recombination and 20 with *sph-glp* recombination (Fig. 4e, Fig. S40b, Fig. S42), yielding 15 new derivatives, with product generation probabilities of 78.9% and 70%, respectively. (Fig. S40, Figs. S42 and S43). Recombinants produced diverse products, with specific variants achieving highest yields for each: R5 (F1) for **Rp1**, R15 (F11) for **Rp3**, R16 (F12) for **Rp4**, R21 (F17) for **Rp6**, R28 (Fs5) for **Rp7**, R32 (Fs9) for **Rp8**, R35-36 (Fs12/Fs13) for **Rp9**, R38 (Fs15) for **Rp10**, and R42 (Fs19) for **Rp12** (Fig. 4e, Fig. S40, Figs. S42 and S43). Overall, the recombination sites distributed widely across C, A and T domains, with optimal results in the A domain’s A6-A10 motif, C domain’s C2/C5/C6 motifs, and T domain.

Based on the characteristics of the RACE method, the most crucial factor in rational modification is to restrict the recombination region to ensure the fusion site is located at the target position. We thus have developed two strategies for the rational design of target sites. One strategy is based on the *de novo* design, which identifies the optimal fusion sites upstream and downstream by using specific modules through the decomposition of the gene cluster, and ultimately selects these sites for rational modification (Fig. S29). The other strategy involves using a resistance gene to restrict the recombination region in gene clusters and conducting two rounds of RACE to identify the optimal fusion sites (Fig. S44). In the following section we comprehensively employed both strategies to identify the optimal upstream and downstream fusion sites. 1) To modify rzmA’s second residue (change L-Tyr to L-Gln from *glp*), we selected fusion point F2 from recombinant D8 (Fig. 5a and Fig. S45a). Most recombinants produced the target compound via one round of RACE (except Fs3). The optimal downstream fusions Fs1 (R45) and Fs6 were identified (R50) (Figs. S45 and S47), finalizing the product Rp14 (Fig. 5a and Fig. S48); 2) For sph’s third residue (L-Ser replacement with L-Gln from *glp* modules) (Fig. 5b, Fig11b and Fig. S46), two rounds of RACE yielded the combination of optimal fusions: F12 (R69, first round) and Fs11/Fs12 (R83/R84, second round), finally creating **Rp17** and **Rp17-2** (Fig. 5b, Fig. 11b and Figs. S46, S47 and S50); 3) Replacing *sph* modules with *gdn* modules 5-8 (Fig. 5c, Fig. 11b and Fig. S39a, c) generated six derivatives (**Rp19** series). First-round RACE identified F4 (R88) as optimal, which was then combined with second-round fusions Fs4 (R113), Fs15 (R124), Fs22(R131), Fs23 (R132) and Fs27 (R136) to design the final products **Rp19, Rp19-2, Rp19-3, Rp19-4/Rp19-5**, and **Rp19-6**, respectively (Fig. 5c, Fig. S11b and Fig. S49b and S51). 4) Engineering *sph* with *gdn* modules 10-11 via two rounds of RACE created Rp21 and Rp21-2 (Fig. 5d, Fig. S52). The optimal first-round fusion F1 (from R139) was combined with second-round fusion Fs7 to generate **Rp21**, and with Fs23 to produce **Rp21-2** (Fig. 5d, Figs. S53-S55).

**Fig. 5.**
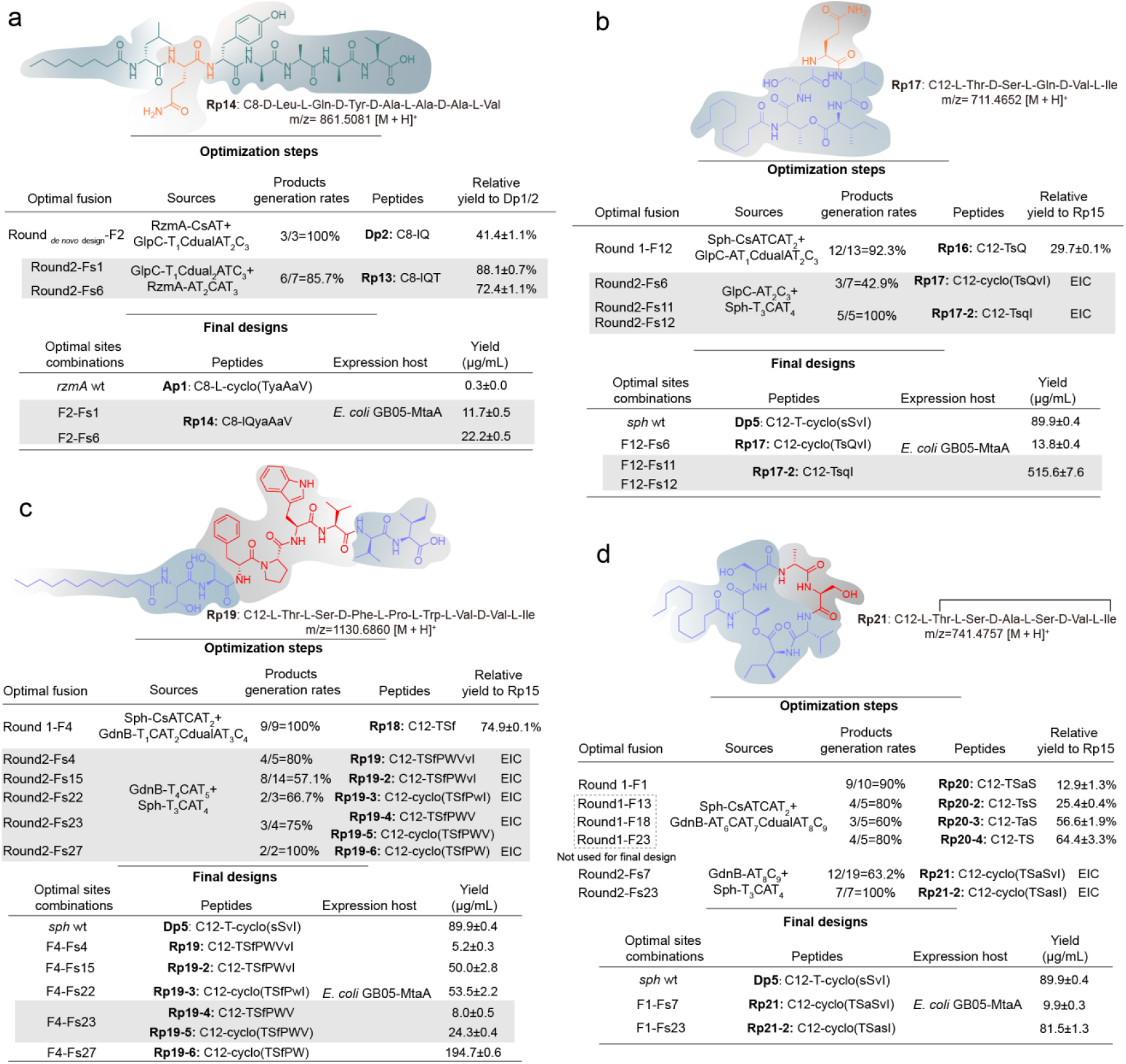
Statistical analysis of rational design. **a**. summary of Rational Design to create product **Rp14**, this process consists of the optimization step followed by a final design step. The optimization focuses on selecting optimal upstream and downstream fusion sites through RACE and recombinant quantification. The final step applies the selected sites (optimal fusion) to design the full-length gene cluster. **b**. summary of Rational Design to create product **Rp17. c**. summary of Rational Design to create product **Rp19. d**. summary of Rational Design to create product **Rp21**.

In summary, the median relative yields of recombinants producing **Rp13, Rp16, Rp18**, and **Rp20** under the rational design strategy were 36%, 21%, 62% and 12%, respectively, with an overall median relative yield of 21%. These results illustrate the substantial variability in pairing efficiency across different modules. We propose two rational design strategies to constrain recombination within defined target regions, thereby enabling site-specific optimization. 1) The first strategy adopts a “building block” framework, wherein the gene cluster is divided into discrete modules or domains for point-to-point optimization (Fig. S29). This approach is applicable to both *de novo* design and rational design, it supports simultaneous optimization across multiple sites, after which the best-fitting variant at each site can be selected for integration into the final construct. For non-initiating modules, an additional lipid-chain addition (using a Cs domain containing module) step is required to enable quantitative analysis of recombination events. 2) The second strategy utilizes resistance markers to define recombination regions, and the final construction is achieved via RedEx-mediated seamless editing (Fig. S44). This method is particularly well-suited for site-directed and rational design applications. Collectively, these two approaches provide generalizable solutions for RACE-based rational engineering.

#### Analysis the fusion sites governing NRPS engineering

The RACE generated an extensive recombination dataset including 830 recombinants and 284 classes of NRP derivatives, offering valuable insights into NRPS engineering rules by detailed analysis of the link of recombination fusion sites to their desired products. We firstly investigated the overall frequency of recombination sites to discover recombination hotspots. For consistency, all domains and motifs were annotated using antiSMASH 7.0 [57]. The fusion sites are distributed across the C-A-T module, most of located in the A (323 recombinants) and C (223 recombinants) domains as hotspots (Fig. S56a). However, normalizing by domain sequence length (fusion frequency per unit length) shows the T domain and A-T junction as more prominent hotspots (Fig. S56b). The independent analysis of the recombinants of individual gene clusters showed similar trends (Fig. S56c-f). The specific recombination hotspots in each domain were also identified, the c6/7, c5, c4, and c2 motifs were predominant in the C domain with c6/7 being the most frequent, and the hotspots clustered near motifs a2, a3, a6, and a8 in the A domain (Fig. S56g, G1-G2 groups).

We next evaluated each recombination sites by using the success ratio (Productivity=number of productive fusion sites/total fusion sites×100%) to discover the most productive regions for generating detectable products (Fig. S56h). The statistics of product-generation probabilities across all recombinant groups revealed both high-efficiency (82.2%) and putative low-efficiency (58.3%) recombination regions (Fig. 6a). Overall, the median relative yield was 24% across 430 groups, with 22% of recombinants exhibiting relative yields above 100% and 37% exceeding 50% (Fig. 6b). The sites located in the c4 and c5 motifs of C domains (Sites S1-S6; Figs. 6c, d), and the a8 and a10 motifs of A domain (Sites S7-S13) are the most productive hotspots in corresponding domains, with the success ratio reaching 82%, 86%, 72% and 79%, respectively. Productive sites were also identified in the T domain (success ratio 78%, Sites S14-S16; Figs. 6c, d), containing the previously documented XUT site (S15) obtained by bioinformatics-guided evolution analysis[15], which demonstrate that the experimental RACE result fits well to the bioinformatics analysis. The overall scatter plot reveals a clear peak in the a8-T region, highlighting its advantage as a high-efficiency recombination region. Sites S8/S9, S12/13, and S15 (XUT) also appear to be optimal fusion sites based on recombination frequency and success rate (Fig. 6a). However, for different gene clusters, their productive recombinant hotspots are not identical. For example, in *rzmA* and *holA* evolution, hotspots occurred at S1-1 and S1-2 (C domain), S12 (A domain), and S1-3 (T domain) (Fig. S57), whereas *glp* evolution showed hotspots at S4 (C domain), S3-1 (A domain), and S3-2 (T domain) (Fig. S59). The most productive sites for each cluster are summarized in Table S3 and Figs. S57-S61. The pronounced differences in fusion sites observed across distinct gene clusters and even among individual modules underscore the necessity of customized synthesis.

**Fig. 6.**
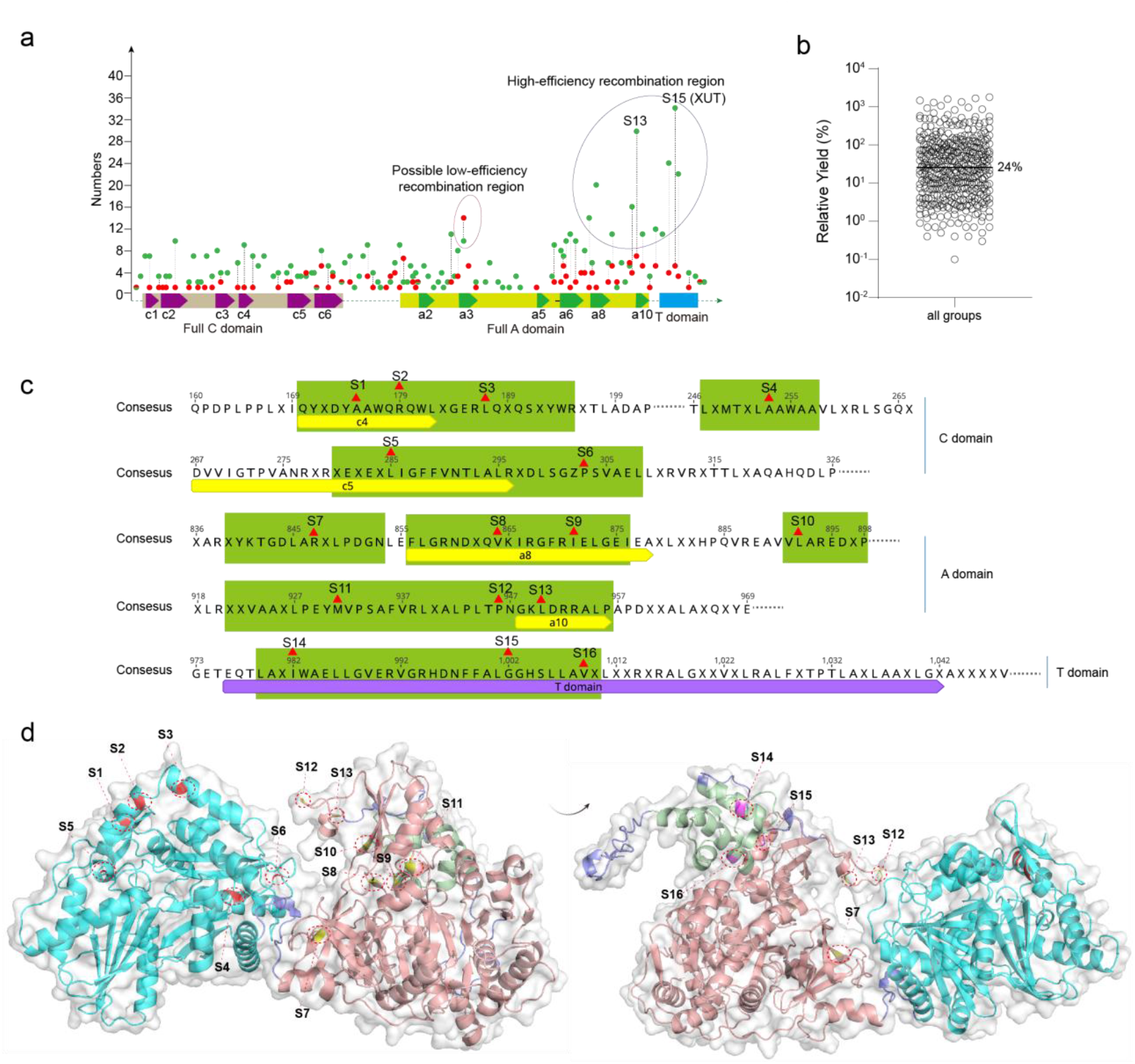
Statistical analysis of fusion sites for all groups. **a**. summarizes all recombination sites across both rational and irrational design sections. Groups yielding products are shown in green, while those without detectable products are shown in red. Fusions involving identical modules are counted only once. **b**. relative yields and their median values across all groups. For groups producing multiple products, relative yields represent the sum of the yields of all final products. **c**. given the widespread distribution of recombination sites across the CAT region, only the top-ranked fusion sites (based on productivity) are shown here, marked by red triangles for specific sites and green squares for hotspot regions in the consensus sequence. The consensus sequence (50% identity threshold) was generated from the multiple sequence alignment of CAT modules, including RzmA-CAT2, RzmA-CAT3, HolA-CAT2, GdnB-CAT4, GdnB-CAT7, GlpC-CAT3, GlpC-CAT5, and IcoA-CAT4. **d**. presents the predicted protein structure of the RzmA-CAT3 module, generated using AlphaFold2, to visualize the spatial positions of the identified fusion sites.

The issue of C-domain substrate specificity for exchange of module selection is also discussed in the Supplementary Materials Data S1 (Analysis of Recombination section). Most C domains lacked strict donor specificity, with many recombinants producing derivatives despite deviating from presumed donor preferences (Table S2, G1 groups).

The recombination occurring at the linker regions of domains is rare but still productive, and the S6-1(near reported XU site) in C-A linker, S6-2 in A-T linker, and S6-3 in T-C linker were identified as promising fusion sites in these junctions (Fig. S62). The broad distribution of functional recombination sites aligns well with bioinformatic predictions and covers nearly all previously reported fusion sites[15, 16, 21, 24, 58-60]. The variation in evolutionary outcomes across gene clusters exhibits the diversity and complexity of natural NRPS evolution (Figs. S56-S61), reinforcing that recombination-driven evolution is a powerful and effective strategy for NRPS engineering.

Furthermore, analysis of non-productive recombination events revealed that the a2 and a3 motifs are frequent recombination hotspots in the A domain, but they yield functional products at remarkedly lower rates than the a6, a8, and a10 motifs (Figs. S14g, h). This may be due to the presence of highly conserved residues in a2 and a3 motifs, which are critical for substrate specificity or structural interactions[49, 58, 61]. These essential residues are likely incompatible with the distantly related gene cluster, making the regions unsuitable for productive recombination.

We comprehensively cataloged all fusion sites identified through high-throughput recombination and rational design. Recombination fusion sites were defined as a single locus if they had fewer than five amino acid differences. Detailed analyses are available in the Supplementary Materials Data S1(Analysis of Recombination section; Fig. SD1). These results illustrate the high variability in optimal fusion site selection across different modules, arguing against the feasibility of a universal site for all NRPS engineering. Using multiple fusion sites for given design may improve success rates of chimeric assembly lines. Notably, productive sites for a given module exchange often yielded comparable levels of product, suggesting that the final production is mainly determined by the compatibility between the paired modules (Figs. S5, S8, S13-1, S20-2, S30, S37, S47, S50, S51, S55). These findings demonstrate module selection and target site customization are crucial for effective NRPS engineering. Tuning these considerations to target gene clusters will be essential to circumvent engineering failure and achieve efficient, customized biosynthesis.

#### Unusual Bis-lipopeptides and their bioactivity

Early studies have demonstrated that the lipid chain of lipopeptides is critical for their biological activity[62, 63]. For instance, a class of chemically synthesized bis-lipopeptides (polymyxin analogues) exhibited at least a 50-fold increase in antibacterial activity against polymyxin-resistant *Pseudomonas aeruginosa*[54]. Based on this insight, we aimed to introduce a second fatty-acid chain into the previously reported lipopeptide C8-rzmA[19] by introducing the unique IcoA C3-Cs4 module, thereby enabling the high-throughput artificial construction of bis-lipopeptides (Fig. 7a). Using both inter-gene cluster evolution and continuous recombination strategies, we generated 170 distinct recombinants, yielding 55 classes of bis-lipopeptides encompassing 181 newly identified structures, with relative yields ranging from 1.5% to 602.2%. High-yielding variants were obtained using the T_1_C_dual_AT_2_C_3_Cs_4_AT_3_ module paired with fusion-site sets such as S16/S6-2 (upstream/downstream sites), S13/S14, S3-2/S3-2, and S1-3/S14, as well as using the T_2_C_3_Cs_4_AT_3_ module together with fusion site including S1-3/Sa31, S15/S15, and S15/S1-3 (Fig. 7b, c). These findings demonstrate the high efficiency of the RACE strategy and establish a foundation for the future customized synthesis of unusual bis-lipopeptides.

**Fig. 7.**
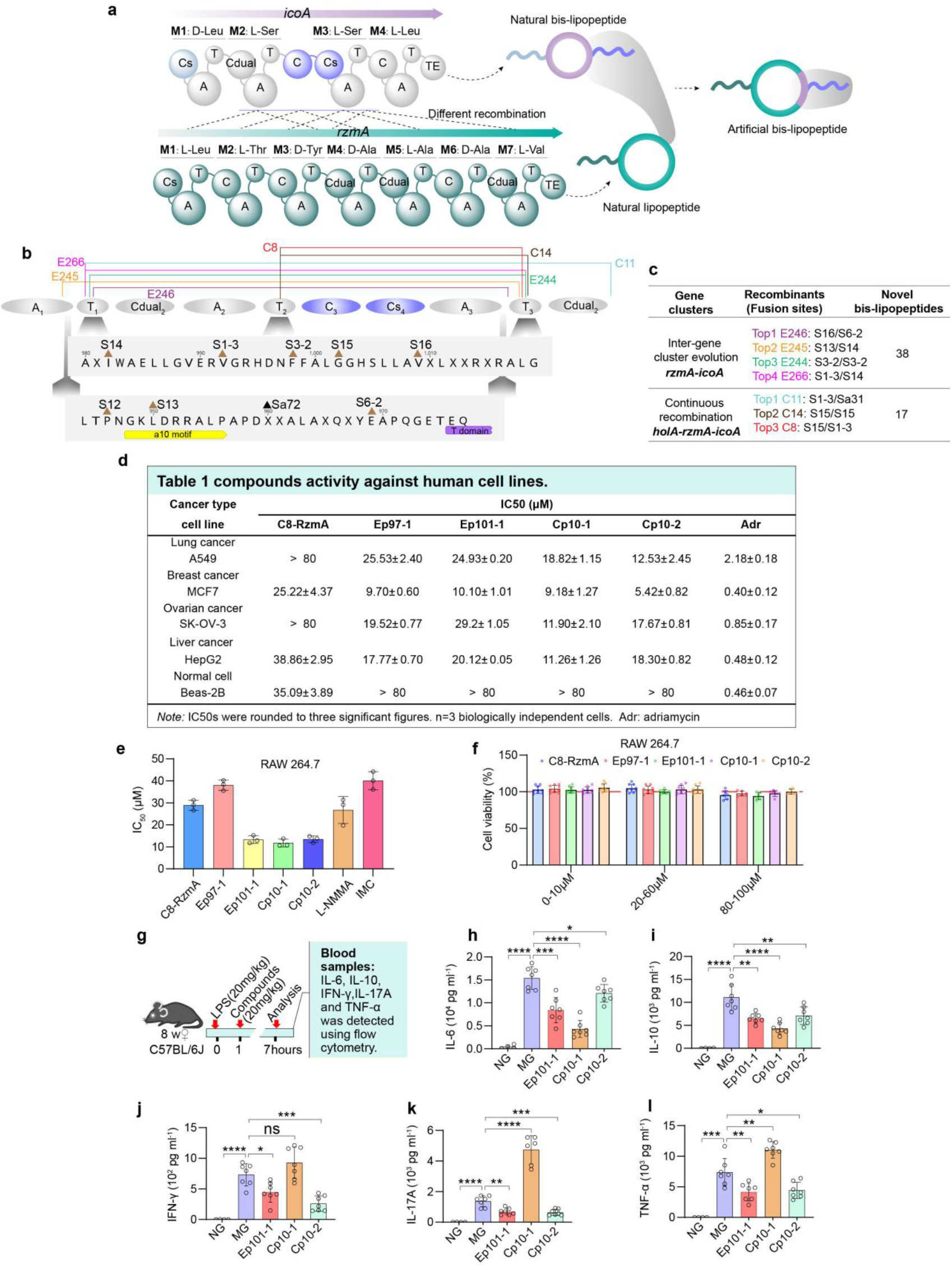
(next page). Evaluation of the anti-tumor and anti-inflammatory activities of compounds. **a**. artificial construction of bis-lipopeptides by C-Cs module recombination using the race strategy. **b** and **c**. analysis of the fusion sites in the top-yielding recombinants, including E244, E245, E246, and E266 obtained from inter-cluster evolution, and C8, C11, and C11 derived from continuous recombination. **d**. cells were treated with compounds (**C8-RzmA, Ep97-1, Ep101-1, Cp10-1**, and **Cp10-2**) at various concentrations to determine IC_50_ values. Data are expressed as mean±s.d. (n=3). Adr: adriamycin. **e**. the RAW264.7 cell line was exposed to compounds alongside positive controls (L-NMMA and IMC) at different concentrations to calculate IC50 values. Data are presented as mean±s.d. (n=3). L-NMMA: NG-monomethyl-L-arginine monoacetate salt (nitric oxide synthase inhibitor); IMC: indomethacin. **f**. the effects of compounds on cell viability were assessed using the murine macrophage cell line RAW264.7. Data are reported as mean±s.d. (n≥6). **g-l**. anti-inflammatory activity of compounds evaluated using an acute inflammation model. **g**. schematic illustration of the protocol for administering LPS and/or compounds in C57BL/6J mice. **h-l**. effects of compounds on the production of inflammatory cytokines in LPS-stimulated mice: interleukin-6 (IL-6, **h**), interleukin-10 (IL-10, **i**), interferon-γ (IFN-γ, **j**), interleukin-17A (IL-17A, **k**), and tumor necrosis factor-α (TNF-α, **l**). Inflammatory mice were treated with compounds at a dosage of 20 mg kg^−1^ (10 mpk). **h-l**. data are presented as mean ± s.d. (NG, n=4; MG, n=7). NG: negative group (treated with saline only); MG: model group (treated with LPS only). Statistical analysis was performed using a two-tailed Student’s t-test. ns: not significant; *p < 0.05; **p < 0.01; ***p < 0.001; ****p < 0.0001.

From these recombinants, we isolated four novel bis-lipopeptides (**Ep97-1, Ep101-1, Cp10-1, Cp10-2**; Fig. SD2), whose structures were elucidated by NMR and Marfey’s analysis (Figs. SD3-SD25, Tables S8 to S11), followed by evaluation of their bioactivities. All showed moderate antitumor activity with improved tumor selectivity over the original product C8-RzmA[19] and excellent anti-inflammatory effects *in vitro* and *in vivo*[64] (Fig. 7d-l and Fig. SD2). **Cp10-1** and **Cp10-2**, differing only in ring size, displayed distinct cytokine modulation, implying structure-dependent mechanisms. Detailed analyses are available in the Supplementary Materials Data S1(Evaluation activities of derivatives section).

## Conclusions

Recombination-driven evolution increases the diversities of NRPSs in nature, and provides a powerful biological paradigm and an ideal blueprint for NRPS engineering. By emulating the core mechanisms of HGT and homologous recombination in natural evolution, we developed the RACE strategy, which enables high-throughput modification of NRPSs within a remarkably short timescale. Building on this foundation, we further established the rational strategy based on RACE that achieves precise engineering operations, including *de novo* design and site-specific modification of NRPSs. Comprehensive analysis of large-scale recombination data revealed two key determinants for customized NRPS biosynthesis: (i) identification of compatible pairing modules for target modules/domains, and (ii) optimization of fusion sites for target modules/domains. Both challenges can be effectively addressed through RACE, providing a broadly applicable framework for tailored NRPS design. Furthermore, analysis of recombination sites and success ratio revealed numerous recombination events distributed across the C-A-T module, providing an extensive landscape for recombination-driven evolution of NRPSs. Among them, a high-efficiency recombination region (a8-T region) was identified, along with several highly promising sites suitable for rational modification, such as the newly discovered S8/S9 (XQVKIRGFRIEL, near the reported A8b and A8c sites[23]), S12/13 (TPNGKLDRR, near the reported site of A10 motif [23]), and S15 (ALGGHSLL, as previously reported as XUT[15]). These sites constitute a valuable resource for precision NRPS engineering. It is noteworthy that these sites may be applicable only to specific gene clusters, as differences in evolutionary trajectories among different gene clusters could restrict the universality of such “plug-and-play” recombination sites.

Over the past decades, substantial effort has been devoted to dissecting the interface interactions of modular NRPSs and PKSs from the crystallization perspective[65, 66]. Guided by these structural insights, researchers have sought to define minimal-disruption cleavage sites for reprogramming, yielding numerous engineering successes[16, 21, 24, 67]. From an evolutionary standpoint, such structure-based definitions of fusion points appear misaligned with the mechanisms that drive natural NRPS diversification, which appear driven less by the fine-tuning of protein-protein interfaces than by DNA-level recombination, privileging stretches of sequence homology between gene clusters over the precise positioning of cleavage sites. Strikingly, recent work from the groups of Bode and Piel has harnessed co-evolutionary analyses of NRPS and *trans*-AT PKS gene clusters to identify potential recombination fusion sites, enabling successfully engineering design[15, 40]. These advances show the notion that the programmability of modular NRPSs and PKSs can be maximized by following the rules “imprinted by evolution”[68]. Unlike conventional combinatorial biosynthesis strategies centered on predefined fusion sites, we established a recombination-driven engineering approach, RACE, that directly exploits recombination to enable controllable NRPS evolution. The fusion sites generated during this process can further guide “site-directed engineering strategies”. Accordingly, we propose a conceptual framework of “recombination-directed NRPS evolution”, under which the application of RACE holds strong potential for the artificial customization of bioactive peptides.

The RACE method developed in this work thus has enormous potential to deliberately design unnatural natural products at scale and make them amenable for high-throughput analysis of bioactivities and pharmaceutical properties, eventually making the system a potential game-changer in the discovery and development of potential drug candidates.

## Materials and methods

### Strains, media and growth conditions

For the recombinants which expressed in the host DSM7029 or its engineered derivatives (e.g., DT8), A 500 μL aliquot of an overnight culture was inoculated into 50 mL of fresh CYMG liquid medium supplemented with appropriate antibiotics in a 250 mL flask. The culture was then incubated in a 30°C orbital shaker at 200 rpm for 5 days. On day 4, 1 mL of XAD-16 resin was added. Cells and resin were harvested the following day. Metabolites were eluted by adding 40 mL of methanol. After centrifugation, the solvent was evaporated under reduced pressure using a rotary evaporator. The residue was dissolved in 1 mL of methanol for LC-MS analysis. For recombinants expressed in the host *E. coli* strain, The identical procedure was followed, except that LB medium was used.

### Nomenclature of recombinants, mutants, and products

Plasmid nomenclature was defined using a gene cluster-recombination site format. For example, recombinant plasmid E10 was generated by recombination between the *holA* gene cluster donor and the *rzmA* recipient gene cluster located on the p15A-apra-phiC31 backbone. The upstream recombination junction occurred between residue E4716 in the A_5_ domain of the *rzmA* gene cluster and residue E493 in the A_1_ domain of the *holA* gene cluster, whereas the downstream junction occurred between residue D5415 in the C6 domain of *holA* and residue D5412 in the C6 domain of *rzmA*. Accordingly, this plasmid was designated as p15A-apra-phiC31-km-rzmA-A_5_E4716-holA-A_1_E493-holA-C_6_D5415-rzmA-C_6_D5412.

Recombinant plasmids generated through “Intra-gene cluster evolution were designated” with the prefix A, and their corresponding products were designated **Ap**. Recombinant plasmids arising from “Inter-gene cluster evolution” were designated E, with products designated **Ep**. Plasmids generated via “Continuous recombination” were designated C, with products designated **Cp**. Plasmids generated through “*De novo* design” were designated D, with products designated **Dp**. Plasmids generated through “Rational modification of NRPSs” were designated R, with products designated **Rp**. Recombinant plasmids generated in “Testing RACE for the evolution of distantly related gene clusters” were designated T.

Wild-type products of the *rzmA* gene cluster were designated as follows: C8-Rhizomide A as **Ap1**, and Rhizomide A (C2) as **Ap2**. The wild-type product of the *holA* gene cluster, holrhizin A, was designated **Ap23**. Wild-type products of the glidonin gene cluster (*gdn*) were designated **Ap37-1** (glidonin A), **Ap37-2** (glidonin G), **Ap37-3** (glidonin I), and **Ap37-4** (glidonin L). The wild-type product of the glidopeptin gene cluster (*glp*), glidopeptin B, was designated **Ap79**. The wild-type product of the icosalide gene cluster (*ico*), icosalide A1, was designated **Ep88**, and the wild-type product of the stephensiolide gene cluster (*sph*), stephensiolide I, was designated **Dp6**.

Mutant strains were designated using the format strain::plasmid. For example, the mutant generated by introducing plasmid E10 into strain DSM7029-DT8 for product expression was designated 7029-DT8::E10.

### Mechanistic study of RACE-mediated recombination

PCR-amplified *amp* fragments carrying PMHAs were obtained from the pR6K-amp-ccdB template and recombined with the PCR-derived *p15A-cm* vector using LLHR, to construct H1-H12. Recombinants were selected based on dual antibiotic resistance (ampicillin and chloramphenicol). The recovered culture was plated at various dilution ratios, and the number of individual colonies was counted to calculate the total number of clones based on the dilution factor. Plasmids were extracted from 24 randomly selected colonies and subjected to Sanger sequencing. Sequencing results were used to determine recombination accuracy. Four experimental groups were established: (1) Two pairing regions with one point mutation; (2) Two pairing regions with two-point mutations; (3) Two pairing regions with three-point mutations; and (4) Three pairing regions with two-point mutations. Detailed experimental procedures are provided in tables S4-S7.

### Intra-gene cluster evolution

The rhizomide (*rzmA*), holrhizin (*holA*), glidonin (*gdn*) and glidopeptin (*glp*) gene clusters were conducted intra-gene cluster evolution via RACE. For the *rzmA* and *holA* gene clusters, the toxic gene cassette *amp-ccdB* was inserted into related plasmids to create the gaps for RACE. For the *rzmA* gene cluster, A1, A2 and the uncorrected recombinants on the step that adding the *amp-ccdB* were digested by PmeI and mixed to conducted RACE in *E. coli* GB05-MtaA::pSC101-tet-PBAD-ETγA to constructed recombinants A3 to A43. These recombinants were expressed and quantified using LC-MS and the **Ap1**: C8-Rhizomide A served as the reference (100%) for quantifying the recombination products of *rzmA*. Totally 20 new derivatives were gained including **Ap3**-**Ap22**; For the *holA* gene cluster, A44, A45 and the uncorrected recombinants on the step that adding the *amp-ccdB* were digested by PmeI and mixed to conducted RACE in *E. coli* GB05-MtaA::pSC101-tet-PBAD-ETγA to constructed recombinants A46 to A74. These recombinants were expressed and quantified using LC-MS and the **Ap23**: Holrhizin A served as the reference (100%) for quantifying the recombination products of *rzmA*. Totally 13 new derivatives were gained including **Ap24**-**Ap36**; For the *gdn* gene cluster, A75, A76 plasmids which contained the *cm-ccdB* cassette were digested by PacI and mixed to conducted RACE in *E. coli* GB05-dir to constructed recombinants from A77 to A142. These recombinants were expressed in host 7029-ΔBPN and quantified using LC-MS and the **Ap37-1**: Glidonin A served as the reference (100%) for quantifying the recombination products of *gdn*. Totally 37 novel derivative compound groups were gained including **Ap38**-**Ap51, Ap53-Ap58, Ap60-Ap64, Ap66-Ap76, Ap78**; For the *glp* gene cluster, A144, A145 plasmids which contained the *amp-ccdB* cassette were digested by PacI and mixed to conduct RACE in *E. coli* GB05-dir to constructed recombinants from A146 to A227. These recombinants were expressed in host 7029-ΔBPN and quantified using LC-MS and the **Ap79**: Glidopeptin B served as the reference (100%) for quantifying the recombination products of *glp*. Totally 25 novel derivatives were gained including **Ap81-Ap97, Ap99-Ap102, Ap104-Ap107**. Full detailed information regarding primers, plasmid/mutant construction, and product analysis is provided in the tables S4-S7.

### Inter-gene cluster evolution

The *rzmA* gene cluster served as the acceptor for RACE with *holA* as the donor. A *cm-tetR-tetO-ccdB* cassette was inserted into the A4 domain region of the *holA* cluster via LCHR to construct plasmid E2. *Ampicillin* resistance cassettes were used to introduce defined gaps at specific sites within the *rzmA* cluster, (between Cs-A_1_, A_2_-T_2_, T_3_-C_dual4_, T_4_-C_dual5_, T_5_-C_6_, and T_6_-C_dual7_) resulting in plasmids E3-E8. Plasmid E2 was linearized by HindIII/HpaI digestion to generate the *holA* donor fragment, which was recombined with PmeI-digested *rzmA* acceptor plasmids (E3-E8) via RACE in GBdir. Recombinant plasmids were verified by AflIII digestion, and those containing unique recombination sites were selected based on distinct restriction patterns. RACE recombination of *rzmA* with *icoA* as donor was performed analogously, with the *cm-tetR-tetO-ccdB* cassette inserted between the special C-Cs didomains of *icoA* to generate plasmid E204. All recombinants underwent scarless marker removal via RedEx recombination, were introduced into strain DT8 for expression, and analyzed by LC-MS for product identification and quantification. Full detailed information regarding primers, plasmid/mutant construction, and product analysis is provided in the tables S4-S7.

### Continuous recombination of NRPS

Similar to inter-gene cluster evolution, the first generation of continuous recombination used the *rzmA* gene cluster as the acceptor and the *holA* gene cluster as the donor, with the recombinase systems Redαβγ and RecET employed to maximize the diversity of initial recombinants. Following AflIII digestion-based screening, ten recombinant clones with distinct fusion sites were randomly selected and pooled to serve as the acceptor population for the second round of recombination. The *icoA* gene cluster was used as the donor, and recombination was carried out using Redαβγ-mediated LCHR without prior linearization, to evaluate whether the LCHR system could also effectively mediate RACE.

The resulting second-generation recombinants were identified through AflIII restriction digestion, and those with distinct digestion patterns were electroporated into the DT8 host strain for expression. LC-MS analysis revealed clear product peaks in several clones, and sequencing of productive recombinants identified approximately 21 unique strains (C1-C21). Among them, 17 recombinants produced novel bis-lipopeptides (**Cp1**-**Cp17**). The detailed construction strategy is provided in tables S4-S7.

### De novo design of artificial NRPs

pRSFDute-rzmA-CsAT-rzmA-TE and pRSFDute-rzmA-CsAT-gdnB-C_13_T_13_TE_1_TE_2_ were expressed in the host *E. coli* BAP1(DE3). The resulting mutants produced the product C8-L-Leu (**Dp1**, m/z=258.2 [M+H] ^+^), which was quantified by LC-MS. In the first round of de novo design using RACE, p15A-apra-phiC31-km-rzmA-CsAT-amp-ccdB-P_13_-cm-gdnB-C_13_A*T_13_TE_1_TE_2_ was digested by PmeI to remove *amp-ccdB*, and then recombined with the fragment glpC-T_1_C_dual2_A_2_T_2_C_3_, amplified from the glidopeptin gene cluster, using LLHR in *E. coli* GB05-dir. Three different recombinants were obtained, namely D7: p15A-apra-phiC31-km-rzmA-T_1_L984-glpC-T_1_L1019-P_13_-cm-gdnB-C_13_A*T_13_TE_1_TE_2_, D8: p15A-apra-phiC31-km-rzmA-T_1_G1024-glpC-T_1_G1059-P_13_-cm-gdnB-C_13_A*T_13_TE_1_TE_2_, and D9: p15A-apra-phiC31-km-rzmA-A_1_R949-glpC-A_1_R970-P_13_-cm-gdnB-C_13_A*T_13_TE_1_TE_2_. These recombinants were then expressed and quantified using LC-MS. The **Dp1**:C8-L-Leu(m/z=258.2[M+H] ^+^) served as the reference (100%) for quantifying the first-round product **Dp2**: C8-D-Leu-L-Gln(m/z=386.2[M+H] ^+^). Subsequently, the optimal fusion D8 was used to construct the second round of design. A PmeI-flanked *amp-ccdB* cassette was inserted into D8 to create D11 by LCHR in *E. coli* GB05-red-gyrA462. After digestion by PmeI, the D11 vector was recombined with the PCR-amplified fragment gdnB-T_4_C_5_A_5_T_5_ by the way of RACE, resulting in the creation of recombinants D12-D19 (tables S4-S7). All recombinants were expressed and quantified using LC-MS, with **Dp2** serving as the reference (100%) for quantifying the second-round product **Dp3**: C8-D-Leu-L-Gln-L-Phe(m/z=533.3[M+H]^+^) (fig. S31). Plasmid D23 was constructed using LLHR in *E. coli* BAP1(DE3), employing the fusion sites F2 and Fs3, along with a recombination site derived from a truncated recombinant A97, as mentioned earlier (Intra-evolution of gene clusters of *gdn*). Subsequently, the absolute quantification of **Dp4** (C8-D-Leu-L-Gln-D-Phe-D-Ala-L-Ser-L-Ala-Put; m/z = 832.5291 [M + H]^+^) was performed using LC-HRMS with a synthesized standard. This product was generated by the mutant *E. coli* GB05-MtaA::D23.

For the design of the terminal module, the A140 recombinant (intra-gene cluster recombination of *gdn*) was recombined with the *sph* gene cluster. D28 was digested using SspI to obtain the fragment sph-A_4_T_4_C_5_A_5_T_5_TE-km-HA, which was then recombined with PacI-digested D26 through RACE in *E. coli* GB05-MtaA::pSC101-tet-PBAD-ETγA. This process resulted in the creation of various recombinants from D29 to D54. Six different derivatives, including **Dp7**: C12-D-Met-L-Ile, m/z =445.3095[M + H]^+^ (produced by D29-D34), **Dp8**: C12-D-Met-D-Ser-L-Ile, m/z =532.3415[M + H]^+^ (produced by D35-D37), **Dp9**: C12-D-Met-L-Ser, m/z = 419.2575[M + H]^+^ (produced by D38-D39), **Dp10**: C12-D-Met-L-Ser-D-Val-L-Ile, m/z =631.4099[M + H]^+^ (produced by D40), **Dp11**: C12-D-Met-L-Ser-L-Ala, m/z =409.2946[M + H]^+^ (produced by D41-D45), **Dp12**: C12-D-Met-L-Ser-D-Ala-L-Ile, m/z =603.3786[M + H]^+^ (produced by D46-D51), and D53-D54 (D52 did not produce the targeted compound), were obtained from a total of 26 recombinants in a single round of RACE. The optimal fusion D35 from the group producing **Dp8**, D50 from the group producing **Dp12**, and D45 from the group producing **Dp11**, were selected to design the final cyclized products. D55, D56, and D57 were thus constructed using the sites from D35, D50, and D45, respectively, to produce compounds **Dp13** (C10-D-Ser-L-Gln-D-Phe-D-Ala-D-Ser-L-Ile, m/z = 806.4659[M + H]^+^)/**Dp13-2**(cyclized product), **Dp14** (C10-D-Ser-L-Gln-D-Phe-D-Ala-L-Ser-D-Ala-L-Ile, m/z = 877.5030[M + H]^+^)/**Dp14-2** (cyclized product), and **Dp15** (C10-D-Ser-L-Gln-D-Phe-D-Ala-L-Ser-L-Ala, m/z = 763.4116[M + H]^+^). Full detailed information regarding primers, plasmid/mutant construction, and product analysis is provided in tables S4-S7.

### Rational modification of NRPS

The start unit design involved constructing the donor plasmid R2, which could supply the start unit from the *glp* gene cluster. This plasmid was then recombined with acceptor plasmids R3 and R4 in *E. coli* GB05-MtaA::pSC101-tet-PBAD-ETγA. As a result, recombined plasmids R5 to R43 were created. Among these, the recombinants *E. coli* GB05-MtaA::R5-R9 produced **Rp1**:C10-D-Ser-L-Gln-D-Tyr-D-Ala-L-Ala-D-Ala-L-Val, m/z =863.4873[M + H]^+^ (R10 did not produce the targeted compound **Rp1**); R11 produced **Rp2**: C10-D-Ser-L-Gln-L-Thr-D-Tyr-D-Ala-L-Ala-D-Ala-L-Val(Ile cyclized with Thr), m/z=946.5245[M + H]^+^;R12 did not produced targeted compound in theory; R13-R15 produced **Rp3**: C10-D-Ser-L-Gln-D-Ala-L-Val, m/z=558.3498[M + H]^+^; R16-R17 produced **Rp4**: C10-L-Ser-D-Tyr-D-Ala-L-Ala-D-Ala-L-Val, m/z=725.4288[M + H]^+^ (R18 and R19 did not produce the targeted compound **Rp4**); R20 produced **Rp5**: C10-D-Ser-D-Ala-L-Ala-D-Ala-L-Val, m/z =572.3654[M + H]^+^; R21-R23 produced **Rp6**: C10-D-Ser-D-Gln-D-Ala-L-Ala-D-Ala-L-Val, m/z=700.4240[M + H]^+^; R24-R28, R30 produced **Rp7**: C10-D-Ser-L-Ser-D-Val-L-Ile, m/z=559.3702[M + H]^+^ (R29 and R31 did not produce the targeted compound **Rp4**); R32 produced cyclized derivate **Rp8**: C10-L-Ser-D-Ser-L-Ser-D-Val-L-Ile(Ile cyclized with L-Ser) and **Rp8-2**: C10-L-Ser-D-Ser-L-Ser-D-Val-L-Ile, m/z=628.3917/646.4022[M + H]^+^ (R33 did not produce the targeted compound **Rp8/Rp8-2**); R34-R36 produced cyclized derivate **Rp9**: C10-D-Ser-L-Gln-D-Ser-L-Ser-D-Val-L-Ile(Ile cyclized with L-Ser) and Rp9-2: C10-D-Ser-L-Gln-D-Ser-L-Ser-D-Val-L-Ile, m/z=756.4502/774.4608[M + H]^+^ (R37 did not produce the targeted compound **Rp9/Rp9-2**); R38 produced **Rp10**: C10-L-Thr-D-Ser-L-Ser-D-Val-L-Ile, m/z=642.4073[M + H]^+^ (R39 and R40 did not produce the targeted compound **Rp10**); R41 produced **Rp11**: C10-D-Ser-L-Gln-D-Val-L-Ile, m/z=600.3967[M + H]^+^; R42 and R43 produced cyclized derivate **Rp12**: C10-D-Ser-D-Gln-L-Ser-D-Val-L-Ile (Ile cyclized with L-Ser) and **Rp12-2**: C10-D-Ser-D-Gln-L-Ser-D-Val-L-Ile m/z=669.4182/687.4288[M + H]^+^; For full detailed information on primers, plasmid/mutant construction, and product analysis, please refer to the tables S4-S7.

**For the rational modification of the second residue L-Tyr in the rhizomide by using the second residue L-Gln from the glidopeptins**, plasmid R11 was recombined with *rzmA-T*_*2*_*C*_*3*_*A*_*3*_*T*_*3*_ in *E. coli* GB05-MtaA::pSC101-tet-PBAD-ETγA. This process created recombinants R45-R51, which produced the compound **Rp13**: C8-D-Leu-L-Gln-L-Tyr, m/z=549.3283 [M + H] ^+^. Among these, the optimal recombinant, *E. coli* GB05-MtaA::R50, was selected for the final design. This led to the creation of **Rp14**: C8-D-Leu-L-Gln-D-Tyr-D-Ala-L-Ala-D-Ala-L-Val, m/z=861.5081 [M + H] ^+^, produced by recombinant R53. **For the modification of the *sph* gene cluster at the L-Ser residue by replacing it with specific modules from the *glp* gene cluster**, two rounds of RACE were conducted. In the first round of RACE, recombinants R58-R66 and R68-R70 produced **Rp16**: C12-L-Thr-D-Ser-L-Gln, m/z=517.3232 [M + H]^+^. In the second round of RACE, recombinants R73, R75, and R78 produced **Rp17**: C12-L-Thr-D-Ser-D-Gln-L-Ile, m/z=711.4652 [M + H] ^+^, and recombinants R80-R84 produced **Rp17-2**: C12-L-Thr-D-Ser-L-Gln-D-Val-L-Ile, m/z=630.4073 [M + H]^+^.The derivatives quantified by used **Rp15**: C8-D-Leu-L-Gln-D-Tyr-D-Ala-L-Ala-D-Ala-L-Val, m/z=861.5081[M + H]^+^ as the control which produced by *E. coli* GB05-MtaA::R56. For the modification of the *sph* gene cluster by replacing with the *gdn* gene cluster (modules 5-8), two rounds of RACE were conducted. In the first round of RACE, recombinants R85-R93 produced **Rp18**: C12-L-Thr-L-Ser-D-Phe, m/z= 536.3331 [M + H] ^+^, the optimal fusion F4 in recombinant R88 was selected for second round design. Other derivates also were produced i.e. the **Rp18-2**: C12-L-Thr-D-Phe, m/z= 449.3010 [M + H]^+^, were produced by R94, R96 and R97 (R95 and R98 did not produced products). The **Rp18-3**: C12-L-Thr-L-Val-D-Phe, m/z= 548.3695 [M + H]^+^ were produced by R99-R101. In the second round of RACE, recombinants R110, R111, R113, R114 (R112 did not produced products) produced **Rp19**: C12-L-Thr-L-Ser-D-Phe-L-Pro-L-Trp-L-Val-D-Val-L-Ile, m/z=1130.6860 [M + H]^+^; R115, R117-R120, R122-R124 (R116, R121, R125-R128 did not produced products) produced **Rp19-2**: C12-L-Thr-L-Ser-D-Phe-L-Pro-L-Trp-D-Val-L-Ile, m/z=1031.6176 [M + H]^+^; R129, R131 (R130 did not produced products) produced **Rp19-3**: C12-L-Thr-L-Ser-D-Phe-L-Pro-D-Trp-L-Ile, m/z=914.5387 [M + H]^+^ (Ile cyclized with L-Thr); R132-R134 (R135 did not produced products) produced **Rp19-4**: C12-L-Thr-L-Ser-D-Phe-L-Pro-L-Trp-L-Val, m/z=918.5336 [M + H]^+^ and **Rp19-5**: C12-L-Thr-L-Ser-D-Phe-L-Pro-L-Trp-L-Val, m/z=900.5230 [M + H]^+^ (Val cyclized with L-Thr); R136 and R137 produced **Rp19-6**: C12-L-Thr-L-Ser-D-Phe-L-Pro-L-Trp, m/z=801.4546 [M + H]+ (Trp cyclized with L-Thr). **For the modification of the *sph* gene cluster by replacing with the *gdn* gene cluster (modules 10-11)**, two rounds of RACE were conducted. In the first round of RACE, recombinants R139-R141 and R143-R148 produced **Rp20**: C12-L-Thr-L-Ser-D-Ala-L-Ser, m/z=547.3338 [M + H]^+^, recombinants R149-R151 and R153 produced **Rp20-2**: C12-L-Thr-D-Ser-L-Ser, m/z=476.2967 [M + H]^+^, recombinants R156-R158 produced **Rp20-3**: C12-L-Thr-D-Ala-L-Ser, m/z=460.3018 [M + H]^+^ and recombinants R159-R163 produced **Rp20-4**: C12-L-Thr-L-Ser, m/z=389.2647 [M + H]^+^, then the fusion F1 in recombinant R139 was selected for the design at the second round of RACE. In the second round of RACE, recombinants R165, R167-R169, R171, R173-R176, and R179-R181 (R166, R170, R172, R177, R178, R182 and R183 did not produced products) produced **Rp21**: C12-L-Thr-L-Ser-D-Ala-L-Ser-D-Val-L-Ile(Ile cyclized with L-Thr), m/z=741.4757 [M + H]^+^; Recombinants R184-R190 produced **Rp21-2**: C12-L-Thr-L-Ser-D-Ala-D-Ser-L-Ile (Ile cyclized with L-Thr), m/z=642.4073 [M + H]^+^.

### Purification of compounds

For compound **Ep97-1**, the extract of 8 L culture of mutant 7029-DT8::E244 was eluted by a normal-phase silica gel column resulting in fraction of CH_2_Cl_2_:MeOH=10:1 which was further purified by semipreparative reversed-phase high-performance liquid chromatography (RP-HPLC) (column: Agilent ZORBAX Stable Bond-C18; 5 μm, 9.4×250 mm; gradient elution: 0-4min, 65% ACN; 4-29 min, 65-95% ACN; 29.1-36 min, 100% ACN; 36.1-40 min, 65% ACN) to afford **Ep97-1** (4 mg, white amorphous solid) at retention time 19.7 min. For compound **Ep101-1**, the extract of 12 L culture of mutant 7029-DT8::E266 was eluted by gradient elution resulting in fraction of CH_2_Cl_2_:MeOH = 15:1 which was further purified by semipreparative RP-HPLC (column: Agilent ZORBAX Stable Bond-C18; 5 μm, 9.4×250 mm; gradient elution: 0-4 min, 62% ACN;4-29 min, 62-87% ACN; 29.1-36 min, 100% CAN; 36.1-40 min, 62% ACN) to obtain **Ep101-1** (2 mg, white amorphous solid) at retention time 19.5 min. For compound **Cp10-1** and **Cp10-2**, the bacterial extract of 8 L culture of mutant 7029-DT8::C11 was purified by semipreparative reversed-phase high-performance liquid chromatography (RP-HPLC) (column: Agilent ZORBAX Stable Bond-C18; 5 μm, 9.4×250 mm; gradient elution: 0-4min, 70% ACN; 4-23 min, 70-93% ACN; 23.1-30 min, 100% ACN; 30.1-34 min, 70% ACN) to afford **Cp10-1** (5 mg, white amorphous solid) and **Cp10-2** (3 mg, white amorphous solid) at retention time 17.5 min and 22.5min. The **Ep97-1, Ep101-1, Cp10-1** and **Cp10-2** were dissolved intoDSMO-*d*_*6*_ for NMR recording (Bruker Avance III-600 spectrometer) and the NMR data were summarized in tables S8-11 and the NMR spectra were shown in figs. SD3-SD25, respectively.

### Compounds characterization

The structure of **Ep97-1, Ep101-1, Cp10-1** and **Cp10-2** were identified by ^1^H, ^13^C, DEPT, COSY, HMBC and HSQC. The compounds data were summarized in tables S8-S11. For **Ep97-1**, HRMS (m/z): [M + H]^+^ calcd. for C_53_H_87_N_8_O_14_, 1059.6337; found, 1059.6325; For **Ep101-1**; HRMS (m/z): [M + H]^+^ calcd. for C_50_H_82_N_7_O_12_, 972.6016; found, 972.6019; For **Cp10-1**, HRMS (m/z): [M + H]^+^ calcd. For C_47_H_83_N_7_O_12_, 938.6173; found, 938.6198; For **Cp10-2**, [M + H]^+^ calcd. for C_47_H_84_N_7_O_12_, 938.6173; found, 938.6196. Compounds were dissolved in 500 μL 6M HCL. Next, the solution was heated at 95°C for 18 h. The resulting hydrolysates of the compounds were dried under vacuum and then re-dissolved in 200 μL ddH_2_O. To derivatize the amino acids, 25 μL 1 M NaHCO_3_ and 200 μL of a 1% Marfey’s reagent solution in acetone were added and mixed. The reaction mixtures were then incubated at 40 °C for 1 h and quenched with 10 μL 2 M HCl. The amino acid standards were derivatized using a similar method. The retention times and molecular weights of the compounds and the standard amino acids are listed in table S11.

### Evaluation activities of derivatives

**Cell culture studies:** A549, MCF-7, SK-OV-3, HepG2, Beas-2B, RAW264.7 cells and their specialized media were obtained from Procell Life Science & Technology Co., Ltd. (Wuhan, China). These cultures were maintained in an incubator at 37 °C in 5% CO_2_. **Cell viability assay**: The viability of A549, MCF-7, SK-OV-3, HepG2 and Beas-2B cells were treated with compounds treatments, 5×10^3^ cells per well were seeded into 96-well plates and treated with various concentrations of compounds for 48 h. Cell viability was determined using the CCK8 assay Kit (Dojindo, CK04). **Inflammation experiments *in vitro* and *in vivo***: Female C57BL/6J mice, aged 8 weeks, were obtained from Shandong Pengyue Laboratory Animal Technology Co., LTD in Shandong, China. Mice were intraperitoneally injected with LPS at a dose of 20mg/kg on the first day. After 24 hours, mice were intraperitoneally injected with the compound at a dose of 20mg/kg. Blood samples were collected after 7 hours for detection of inflammatory cytokines (BD™ Cytometric Bead Array (CBA) Mouse Th1/Th2/Th17 CBA Kit,560485) by flow cytometry. All animal experiments were conducted in accordance with the guidelines formulated by the Ethics Review Committee for Laboratory Animals, School of Life Sciences, Shandong University, with the approval number SYDWLL-2025-131. RAW264.7 cells were seeded into each well of a 96-well plate in 5×10^3^ cells and treated with various concentrations of compounds for 48 h. Cell viability was measured by CCK8 assay kit (Dojindo, CK04). The classic Griess method was used to determine the content of NO released by the compound on LPS-induced RAW264.7 cells. RAW264.7 cells were seeded into each well of a 96-well plate at 5×10^3^ cells, pretreated with the compounds 0.5 hours in time, and incubated with 1ug/ml LPS for 48 hours. The supernatant was detected using the nitric oxide detection kit (Beyotime Biotechnology, S0021S) to determine the release amount of NO.

## Acknowledgments

We would like to thank Zhifeng Li, Guannan Lin, Jingyao Qu, Haiyan Sui and Xiangmei Ren from the Core Facilities for Life and Environmental Sciences, SKLMT of SDU for UPLC-HRMS, NMR and HPLC analysis. This study was supported by the National Natural Science Foundation of China (32201195, 32570094, T2250710184, 32350010, 32170038), the Key R&D Program of Shandong Province, China (2025CXPT191), the Science and Technology Innovation Program of Hunan Province (2025RC3210, 2025RC4019), the 111 Project (B16030), Qingdao Science and Technology Demonstration Projects for the Benefit of the People (25-1-5-xdny-29-nsh), Young Talent Development Program of SKLMT (M2025YA03), and SKLMT Frontiers and Challenges Project (SKLMTFCP-2023-05).

## Data, code, and materials availability

The strains, plasmids, mutants used and constructed in this work and primers used in this work are listed in the supplementary tables and Figures. Correspondence and requests for materials should be addressed to L.Z., Q.T., Y.Z., or X.B.

